# Tumor-on-a-chip platform to interrogate the role of macrophages in tumor progression

**DOI:** 10.1101/2020.05.27.119636

**Authors:** Ye Bi, Venktesh S. Shirure, Ruiyang Liu, Cassandra Cunningham, Li Ding, J. Mark Meacham, S. Peter Goedegebuure, Steven C. George, Ryan C. Fields

## Abstract

Tumor-infiltrating leukocytes, in particular macrophages, play an important role in tumor behavior and clinical outcome. The spectrum of macrophage subtypes ranges from antitumor “M1”-type to protumor “M2”-type macrophages. Tumor-associated macrophages (TAMs) typically display phenotypic features of both M1 and M2, and the population distribution is thought to be dynamic and evolve as the tumor progresses. However, our understanding of how TAMs impact the tumor microenvironment remains limited by the lack of appropriate 3D in vitro models that can capture cell to cell dynamics at high spatial and temporal resolution. Using our recently developed micro-physiological “tumor-on-a-chip” (TOC) device, we present here our findings on the impact of defined macrophage subsets on tumor behavior. The TOC device design contains three adjacent and connected chambers in which both the upper and lower chambers are loaded with tumor cells while the central chamber contains a dynamic, perfused, living microvascular network. Introduction of human pancreatic or colorectal cancer cells together with M1-polorized macrophages significantly inhibited tumor growth and tumor-induced angiogenesis. Protein analysis and antibody-based neutralization studies confirmed that these effects were mediated through production of chemokines CXCL9, CXCL10, and CXCL11. By contrast, M2-macrophages mediated increased tumor cell migration into the vascularized chamber and did not inhibit tumor growth or angiogenesis. In fact, single-cell RNA-sequencing showed that M2 macrophages further segregated endothelial cells into two distinct subsets, corresponding to static cells in vessels versus active cells involved in angiogenesis. The impact of M2 macrophages was mediated mostly by production of MMP7 and ANGPT2. In summary, our data demonstrate the utility of the TOC device to mechanistically probe biological questions in a 3D *in vitro* microenvironment.

**Insight Box:** Macrophages in the tumor microenvironment are key determinants of tumor behavior and clinical outcome. The macrophage subset composition and its functional impact change as tumors progress or during treatment, but adequate models to study this are lacking. We developed a tumor-on-a-chip model of perfused 3D tumor growth to probe the impact of defined macrophage subsets. Our data is consistent with previously described macrophage activity in the tumor microenvironment, and provides potential new molecular targets. Herein, we demonstrate feasibility of probing immuno-oncology questions in a 3D *in vitro* microenvironment and at a spatiotemporal resolution.

## Introduction

Inflammatory cells, such as including macrophages, play a central role in tumor progression, including tumor cell proliferation, angiogenesis, as well as other central processes that shape the tumor microenvironment. Macrophages are particularly interesting as ample evidence has demonstrated a range of phenotypes that can either enhance or inhibit tumor progression depending on the macrophage polarization [1–4]. While classification of macrophages is ongoing, an instructive categorization has been the so-called “M1” and “M2” designations [5, 6]. The former generally refers to macrophages that participate in cell killing, and thus can limit tumor progression, whereas the latter facilitates normal processes such as wound healing and tissue growth and can aid in tumor progression. Understanding the mechanisms that dictate trafficking and differentiation of the macrophage subtypes has generally been limited to 2D cell culture and animal models, both of which have severe limitations. For example, simply plating primary monocytes on a 2D surface initiates differentiation into macrophages, while animal models have limited spatial and temporal resolution to tease apart mechanisms that influence cell trafficking.

We recently previously reported the design of a micro-physiological “tumor-on-a-chip” (TOC) device [7] to study cancer biology. The device design contains multiple adjacent chambers that can be loaded with 3D tissue mimics at different points in time providing tremendous flexibility in the type of tissue and the question(s) that might be addressed. A central feature of the device is a dynamic, perfused, living microvascular network that is ideally suited to examine questions related to angiogenesis in settings such as the tumor microenvironment.

Our current study endeavored to use this device platform to demonstrate its utility to probe questions related to the impact of immune cells (macrophages) on tumor behavior, and specifically interrogate the mechanisms by which macrophages in the tumor microenvironment can modulate tumor cell proliferation and migration, as well as angiogenesis – an important hallmark of cancer [8]. Our results demonstrate that macrophages survive, proliferate, and traffic within the microfluidic device. Furthermore, using a range of biological techniques including single cell RNA-seq, as well as both colorectal cancer (CRC) and pancreatic adenocarcinoma (PDAC) patient-derived cell lines, we discovered that the M1 macrophage inhibited angiogenesis and tumor cell growth through the enhanced production of a series of chemokines (CXCL9, CXCL10, and CXCL11). In contrast, the M2 macrophage stimulated tumor cell migration through the enhanced secretion of a series of proteins that included matrix metalloproteinase 7 (MMP7) and angiopoietin 2 (ANGPT2). These results align with previous reports in the literature regarding the roles of CXCL9, CXCL10, and ANGPT2 [9–12], but, also provide new potential targets (e.g., CXCL11 and MMP7) to limit tumor progression [13, 14]. Further, they demonstrate the utility of the device to probe biological questions in a 3D in vitro microenvironment mechanistically.

## Material and Methods

### Cell lines and reagents

Normal human lung fibroblasts (NHLFs) were obtained from Lonza (Basel, Switzerland), and endothelial colony forming cell-derived endothelial cells (ECFC-ECs) were extracted from cord blood as detailed previously [15]. Both cell types were cultured and used in TOC devices for up to seven passages in complete growth medium from Lonza (FGM-2 for NHLFs and EGM-2 for ECFC-ECs). The CRC cell lines (CRC268, CRC663, CRC1180) and PDAC cell lines (PDAC162, PDAC175) were generated *in vitro* from low passage number patient-derived xenografts (PDXs) established in NOD-SCID mice by the Solid Tumor Tissue Bank and Registry at Washington University in St. Louis. All tumor cell lines were cultured using RPMI1640 medium with 10% fetal bovine serum (FBS) and antibiotics. Monocyte cell lines THP-1 (ATCC^®^ TIB-202™) and U-937(ATCC^®^ CRL-1593.2™), and CRC tumor lines SW480 and SW620 as well as PDAC lines HPAC and Panc1 were obtained from ATCC and cultured in the recommended medium. Reagents used in the *in vitro* macrophage differentiation were as below: Phorbol 12-myristate 13-acetate (PMA), and lipopolysaccharides (LPS) were obtained from Sigma-Aldrich (Darmstadt, Germany). Recombinant human interleukin-4 (IL-4), interleukin-13 (IL-13) and interferon-gamma (IFN-γ) were purchased from PeproTech (Rocky Hill, NJ).

### Cell transduction

The ECFC-ECs were transduced with a green fluorescent protein (GFP)-expressing lentivirus to constitutively express GFP as detailed previously [15]. Colorectal and pancreatic cancer cell lines were transduced using red fluorescent protein (RFP) and puromycin-encoding lentiviral particles obtained from GenTarget Inc. (San Diego, CA) following the recommended protocol. Briefly, cancer cells were seeded into 24-well tissue culture plates and incubated overnight at 37°C in 5% CO_2_. The next day, the medium was replaced with 0.5 ml of complete culture medium and 50ul of lentivirus particles. 72hrs after transduction, the medium was replaced with 1ml complete medium containing 1 μg/ml puromycin for selection to generate stable cell lines expressing RFP.

### Device manufacture, device loading and tissue growth

A schematic overview of the TOC device is presented in Figure 1. The device design and microfabrication have been detailed previously [7]. The microvasculature in the central chamber was created by mixing ECFC-ECs and NHLFs at a 1:2 ratio in fibrin gel (bovine fibrinogen at a final concentration of 10mg/ml and bovine plasma thrombin at a final concentration of 2U/ml). The cells were maintained for seven days in EGM-2 medium to permit development of the microvasculature. Tumor cells were loaded into the side chambers at a concentration of 5 million/ml. When introduced together with immune cells, the tumor cells and immune cells were mixed at a 1:1 ratio with a final cell concentration of 10 million/ml. Devices were kept at 37°C in 5% CO_2_ for another six to nine days and microscope images were taken on days 2, 5 and 9 after loading of the side chambers.

**Figure 1:**
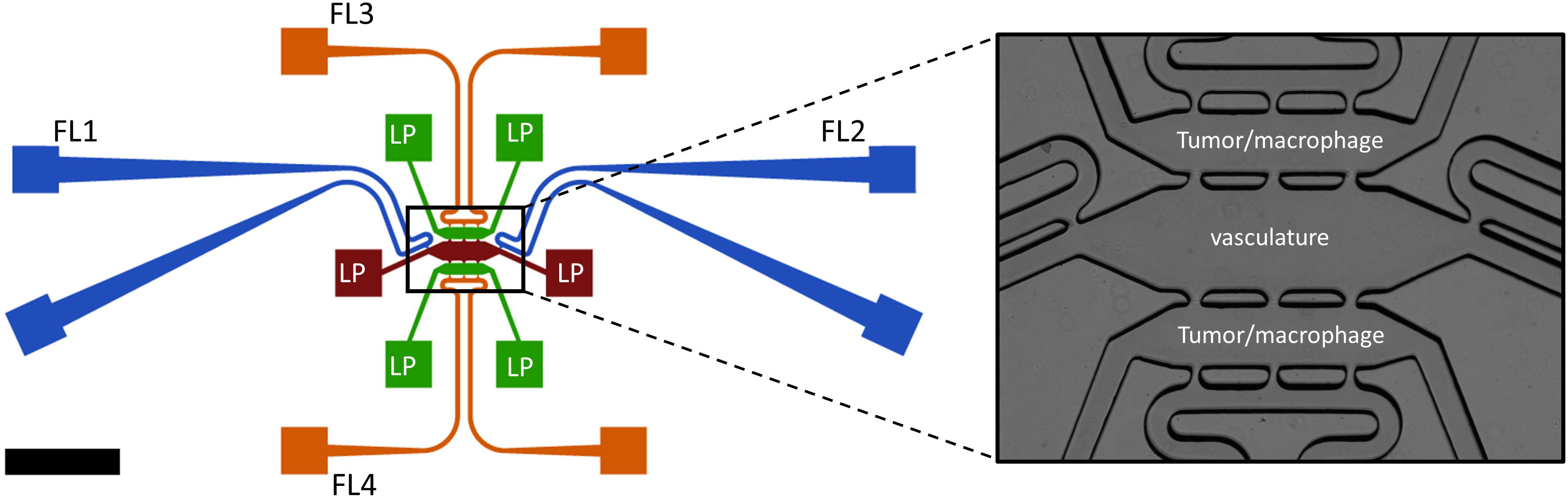
Schematic overview of the three chamber TOC device with tumor-vascular communication. Endothelial cell precursors and fibroblasts are introduced into the central chamber (brown) through one of the connected loading ports (LP). Medium is added through fluidic lines (FL) on both ends of the central chamber (blue), the terminal ends of each fluidic line are connected to a dedicated source and a sink (blue squares). Likewise, the upper and lower side chambers (green) each have loading ports (LP) for introduction of tumor/other cells. The upper and lower chambers also have fluidic lines (green) that can drain excess fluid. The central chamber is connected to the two parallel side chambers through pores that permit both cell and liquid transport (see enlarged image in the right panel).

### *in vitro* activation and differentiation of monocytic THP-1 and U-937 cell lines into macrophages

THP-1 and U-937 cells were cultured at 5×10^5^ cells per well in 6-well plates and activated with PMA at a final concentration of 100 ng/ml for 48 hours. The cells were then washed to remove PMA and any unattached cells. For M1 macrophage differentiation, activated THP-1 and U-937 cells were cultured in 10 pg/ml LPS and 20 ng/ml IFN-γ containing medium for 24 hours. For M2 macrophage differentiation, activated THP-1 or U-937 cells were kept in 20 ng/ml IL-4 and 20 ng/ml IL-13 for 72 hours (Figure S2, [16, 17]). Since M1 and M2 macrophage differentiation required different timeframes, monocytes were activated at different time points to synchronize the harvest of macrophages. Prior to introduction of macrophages into devices, aliquots of cells were stained with antibodies specific for cell markers of M1 and M2 macrophages and analyzed by flow cytometry. The remaining cells were counted and labeled with Violet CellTrace (Thermo Fisher Scientific, Waltham, MA) following the recommended protocol. Subsequently, macrophages were mixed with tumor cells and loaded into devices.

### Flow cytometry analysis

*In-vitro* differentiated THP-1 or U-937 cells were harvested and then stained with antibodies specific for macrophage markers. APC anti-human CD68, APC/Cy7 anti-human Tumor Necrosis Factor alpha (TNF-α), PE anti-human C-X-C motif chemokine 10 (CXCL10), Alexa Fluor 488 anti-human CD206 and PE/Cy7 anti-human Interleukin-10 (IL-10) were used for analysis (all from BioLegend, San Diego, CA). The samples were run on an LSR Fortessa cell analyzer from BD (Becton Dickinson, Franklin Lakes, NJ). For analysis, initial gating was performed on expression of CD68 and positive cells were further analyzed for expression levels of individual markers. Un-activated THP-1 or U-937 cells were used as negative controls.

### Immune fluorescence, imaging and analysis

To perform immune fluorescence (IF) analysis on cells in devices, cells were fixed in 10% formalin for 48 hours. After washing with PBS, cells were permeabilized with 0.2% Triton X-100 in PBS for another 48 hours. Then, primary antibodies diluted in 1% BSA/PBS were added to devices and devices were kept at 4°C for 24 hours. Anti-von Willebrand factor (vWF, at 1:500 dilution), anti-angiopoietin 2 (ANGPT2, at 1:500 dilution) and anti-tissue inhibitor matrix metalloproteinase 1 (TIMP1, at 1:1000 dilution) antibodies were all from Abcam (Branford, CT). After incubation with primary antibodies, devices were washed for 48 hours with PBS. Secondary antibody, goat anti-rabbit conjugated with Alexa Fluor 594 (Thermo Fisher Scientific, Waltham, MA) at 1:500 dilution was added to the devices. After 24 hours at 4°C, devices were washed again for 48 hours, and images were subsequently taken using an EVOS cell imaging system (Thermo Fisher Scientific, Waltham, MA). The vasculature and tumor areas were identified from their respective fluorescent areas in an image using ImageJ 1.52p. Tumor angiogenesis was assessed by measuring the microvessel area (GFP signals) in both side chambers or if no significant blood vessels grown into the side chambers, the ratio of blood vessel areas on day 9 versus day 2 in the central chamber was used as an alternative to show the vasculature density changes. Tumor cell proliferation was assessed by measuring the tumor cells area (RFP signals) in both side chambers and tumor invasion was assessed by measuring the tumor area (RFP signals) in the central chamber at the end of experiments.

### Immunohistochemistry

To perform immunohistochemistry (IHC) on formalin-fixed, paraffin-embedded human tissues, paraffin blocks of the samples were sectioned at 5μm thickness and sections were mounted onto glass slides. After de-paraffinization and rehydration, endogenous peroxidase was blocked with 0.3% H_2_O_2_ in methanol and then antigen retrieval was performed with a heated citrate buffer solution. Anti-CD31 (#JC70A, Agilent, Santa Clara, CA) was used to stain tissue sections. HRP conjugated goat anti rabbit antibody (Abcam, Branford, CT) was then added and finally, DAB-substrate (Vector Laboratories, Burlingame, CA) was added for color development for 2⍰minutes at room temperature. Counter stain was performed with Mayer’s hematoxylin (Thermo Fisher Scientific, Waltham, MA) and cover slips were mounted with Cytoseal XYL (Thermo Fisher Scientific, Waltham, MA).

### Protein analysis and antibody-mediated neutralization

Around 25-40 μl of flow through media was harvested per TOC device containing vasculature + CRC663 + THP-1 M1 or M2 macrophages. Three individual devices were used per condition, and media were pooled. All samples, as well medium controls were subjected to analysis for soluble proteins using the immune-oncology test platform from Olink Proteomics (Uppsala, Sweden). Quality control and NPX values were generated by the company following their standard protocol. For neutralization of specific soluble factors, devices containing vasculature + CRC663 + THP-1 M1 or M2 macrophages were prepared as described above. Antibodies were added to the devices through the feeding medium. Specifically, antibodies to C-X-C motif chemokine (CXCL) 9 (clone 49106 at 0.5 μg/ml), CXCL10 (clone 33036 at 0.5 μg/ml) and CXCL11 (clone 87328 at 0.1 μg/ml) were added to devices with THP-1 M1 macrophages while matrix metalloproteinase 7 (MMP7, clone 111433 at 1 μg/ml) and ANGPT2 (clone 85816 at 1 μg/ml) (all from R&D systems, Minneapolis, MN) neutralizing antibodies were added to devices with THP-1 M2 macrophages. Medium and antibodies were changed every 48 hours.

### Cell harvest and single cell RNA sequencing (sc-RNA seq)

Devices were washed with D-PBS for three times and 20U/ml nattokinase (Japan Bio Science Laboratory Company, JBSL-USA, Walnut Creek, CA) in D-PBS was added to the devices. After 20 minutes incubation at 37°C, cells were harvested from the devices with a pipette and transferred into a 1.5 ml Eppendorf tube. Cells were spun down and resuspended in 500ul of 0.05% trypsin-EDTA, followed by incubation at 37°C for 5 minutes. The cells were washed twice with D-PBS, and re-suspended in 0.04% BSA/PBS. Around 30,000 cells with a viability >80% were analyzed by scRNA-seq at the McDonnell Genome Institute (MGI) at Washington University in St. Louis.

### scRNA-seq data analysis

For scRNA-seq analysis, Cell Ranger (v3.0.2) from 10x Genomics was used for de-multiplexing sequence data into FASTQ files, aligning reads to the human genome (GRCh38), and generating gene-by-cell UMI count matrices. The R package Seurat (v3.1.0) was used for all subsequent analysis [18]. First, a series of quality filters were applied to the data to remove empty droplets/debris and dead cells. To do so, we set the minimum cut-off for number of features and UMI as 200 and 1000, respectively; we set the maximum cut-off for mitochondrial gene expression percentage as 10%. Next, the data were normalized and scaled. Highly variable genes are found for PCA-based feature selection. The cells were then clustered using graph-based clustering (default setting of Seurat). Cell types were assigned to each cluster by manually reviewing the expression of marker genes. The marker genes used were CD34, PECAM1, vWF (endothelial cells); GABARAP, PFDN5, VEGFA (tumor cells); ADM, CTGF, ACTA2 (stroma cells); CD68, CSF1R, MS4A7 (macrophages). Integration was performed with the protocol from the Seurat package. To find differentially-expressed genes between clusters of interest, the function FindMarkers was used with all parameters as default. All the genes returned from the FindMarkers function were treated as differentially expressed genes. For pathway analysis within endothelial cell clusters, cells from each cluster were pulled together for comparison regardless of devices of origin. Genes with absolute log-transformed fold change greater than 0.6 were fed into the Reactome database to perform pathway enrichment analysis. Top enriched pathways are shown.

### Statistics

Statistical analysis was performed using one-way ANOVA tests for multiple comparisons or Student’s t-tests. All data are presented as the mean + standard deviation. Results were considered statistically significant for p<0.05.

## Results

### Tumor cell lines demonstrate growth and heterogenous induction of angiogenesis

The tumor cells lines were loaded into the side chambers of the devices after the microvascular network was formed in the central chamber (Figure 1). Tumor cells were cultured for 9 days and proliferation was assessed. After normalization to the tumor area on day 2, all five early passage tumor lines tested showed a 3-5 folds increase in tumor area by day 9, indicating significant tumor growth (Figure 2A, B). Similar results were seen with the commercial tumor lines (Figure S1). Interestingly, different cell lines displayed diverse invasion capacity over time in the device. CRC663, SW480, and CRC268 tumor lines clearly invaded the vasculature in the central chamber (Figures 2 and S1). These cell lines also induced tumor angiogenesis as new vessels can be observed sprouting from the central chamber into the side (tumor) chambers through the pores between the central and side chambers. Conversely, CRC1180 and the two PDAC lines showed much less invasion and angiogenesis. The level of angiogenesis of tumor lines in TOC devices appeared to correlate with the density of the vascular marker, CD31, in matching parental tumors (Figures 2C, D).

**Figure 2:**
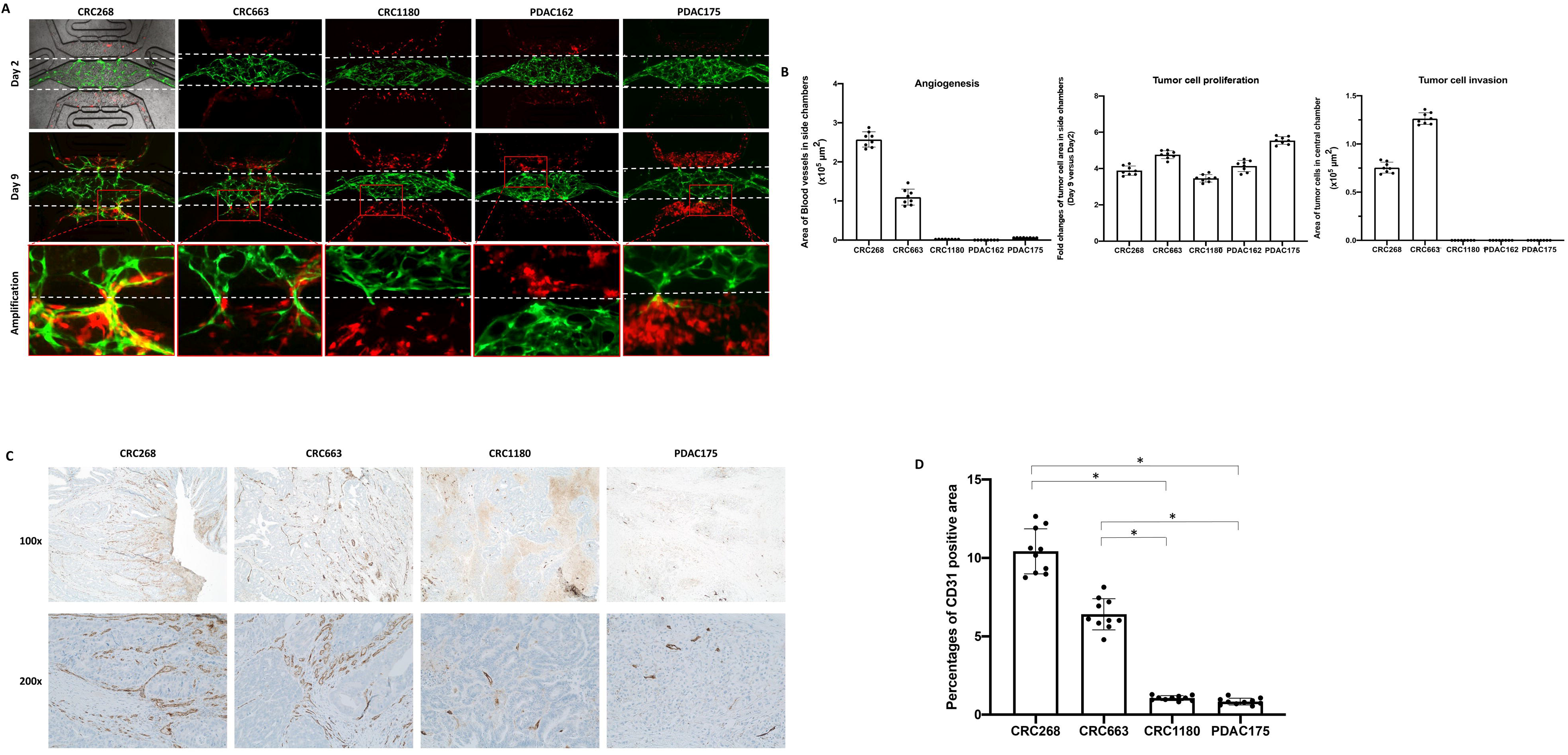
Vascularized TOC devices permit robust growth of colorectal and pancreatic cancer cells. Three colorectal cancer (CRC268, CRC663, and CRC1180) and two pancreatic ductal adenocarcinoma cell lines (PDAC162 and PDAC175) were introduced into the side chambers with pre-formed vasculature in the central chamber. A: Representative immune fluorescence images taken at Days 2 and 9 after tumor cells were introduced. Green: GFP labeled endothelial cells formed vasculature. Red: RFP labelled tumor cells. The white dotted lines indicate the top and bottom boundaries of the central chamber. B: Quantification of angiogenesis, tumor cell proliferation, and migration/invasion on Day 9. Eight devices per tumor cell line were used for quantification. C: IHC images on parental CRC and PDAC tumor sections stained with anti-CD31. Representative images are shown at 100x and 200x magnification. D: Quantification of CD31-positive areas in parental tumors using IHC images. * p<0.05.

### M1 macrophages inhibit tumor invasion, growth, and angiogenesis

To investigate the potential role of macrophages in modulating tumor growth and angiogenesis, we next differentiated monocytic THP-1 cells into M1 or M2 macrophages (Figure S2). Flow cytometry analysis showed that after activation of THP-1 cells with PMA, more than 80% of the cells expressed the common macrophage marker CD68 (data not shown). Differentiation into M1 showed nearly 90% of the cells positive for CXCL10 and among these, approximately 48% cells were also positive for TNF-α (Figure S3A upper panel). Conversely, M2-differentiated macrophages expressed CD206 and IL-10 (Figure S3A, lower panel). Similar differentiation into M1/M2 macrophages was observed using the monocytic cell line, U937 (Figure S3B).

M1 macrophages derived from THP-1 were introduced together with CRC663 or PDAC162 two representative tumor cell lines into the side chambers of vascularized devices. CRC663 represented more aggressive and more angiogenetic tumors while PDAC162 represented relative non-aggressive and non-angiogenetic tumors as shown in Figure 2. Over the course of 9 days, CRC663 showed induction of both angiogenesis and tumor invasion by itself (Figure 3A, first column, 3B). In the presence of un-activated THP-1, there were no detectable differences compared to CRC663 tumor only (Figure 3A, second column). However, in the presence of THP-1 M1, angiogenesis was completely blocked and the microvascular network in the central chamber started to recede by day 9 (Figure 3A, third column, 3B).

**Figure 3:**
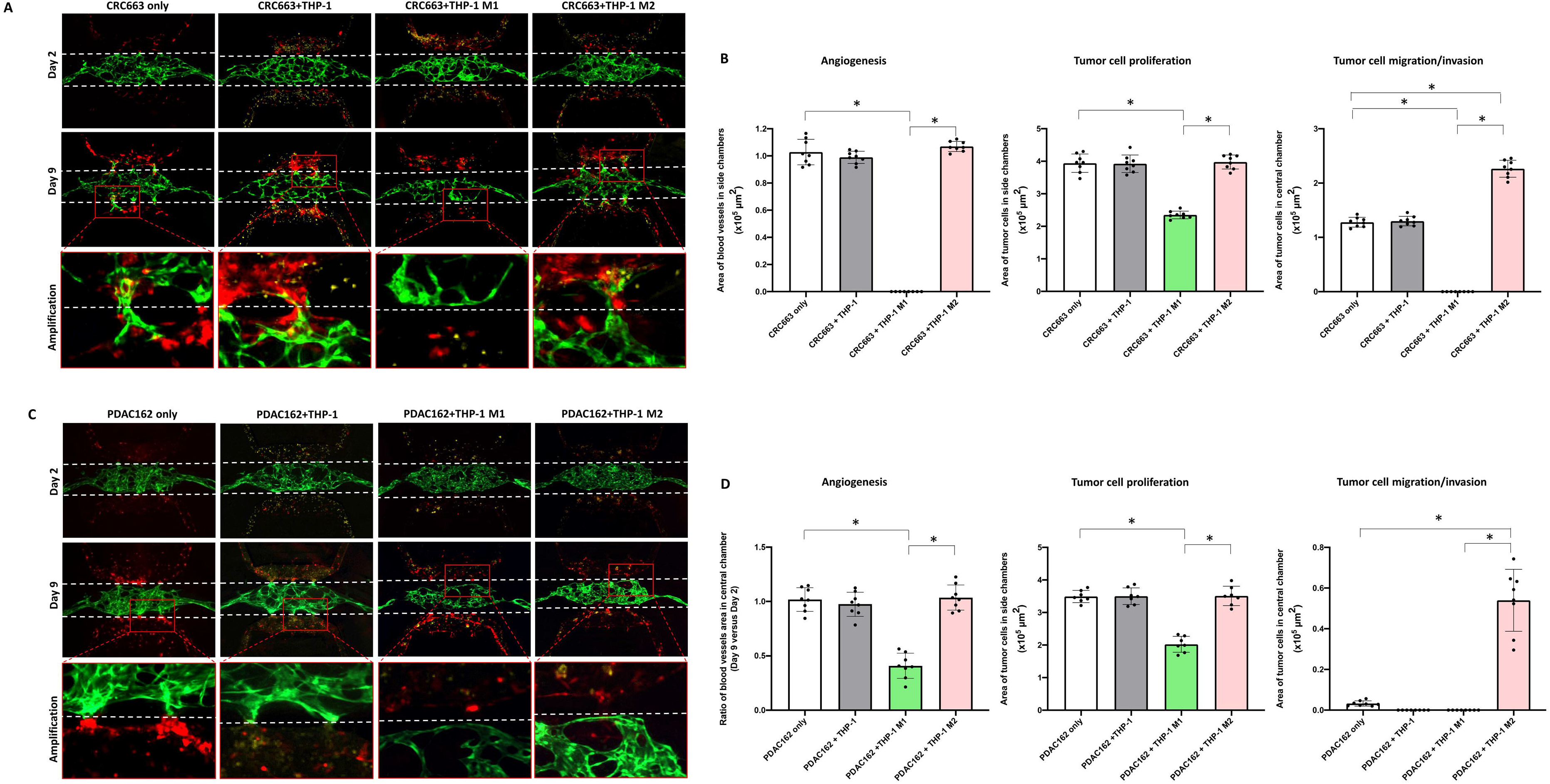
Reconstitution of M1 or M2-differentiated macrophages in TOC devices triggers antitumor and protumor phenotypes, respectively. A, C: Representative immunofluorescence images from Days 2 and 9 after CRC663 (A) or PDAC162 (C) cells with or without THP-1-derived macrophages were loaded into the side chambers. Green: GFP labeled endothelial cells in pre-formed vasculature. Red: RFP labelled tumor cells. Yellow dots: un-activated THP-1 cells or THP-1 derived M1 or M2 macrophages. White dotted line: the top and bottom boundaries of central chamber. Bottom panels represent amplified areas indicated by the red boxes in the middle panels. B, D: Quantification of angiogenesis; tumor cell proliferation, and tumor invasion in the four types of TOC devices per tumor line. Eight devices per group were used for quantification. *: p<0.05.

At the same time, no tumor cells were detected in the central chamber at the end of the experiment (day 9), indicating tumor migration/invasion was also inhibited. Additionally, tumor growth in the side chambers was significantly reduced compared to tumor only devices (Figure 3B). In summary, introduction of M1 macrophages triggered strong anti-tumor effects (inhibition of growth, migration, and angiogenesis) in the devices. Similar phenotypic changes were observed when using PDAC162 tumor cells and THP-1 or THP-1 M1 (Figure 3C, first 3 columns, 3D). U-937 M1 macrophages induced a very similar inhibitory anti-tumor phenotype when cocultured with CRC663 (Figure S4A, first 2 columns, S4B) or PDAC162 (Figure S4C, first 2 columns, S4D).

### M2 macrophages promote tumor cell migration

When M2 macrophages were introduced into devices, alongside tumor cells, tumor growth and angiogenesis were not impacted and were very similar to that observed in devices with tumors only or tumor plus un-activated macrophages (Figure 3A, fourth column, 3B). However, tumor migration into the central chamber was significantly increased. More specifically, the number of CRC663 tumor cells in the central chamber at the end of the experiment (day 9) was two times higher (Figure 3A forth column, 3B). The pro-invasion effect of M2 macrophages on PDAC162 cells was particularly striking. The PDAC162 cells poorly migrated into the central chamber with or without un-activated macrophages, but in the presence of M2 macrophages the number of tumor cells in the central chamber increased by over ten-fold (Figure 3C forth column, 3D). An increase in tumor migration into the central chamber was also observed when M2 macrophages were derived from the monocytic U937 cell line (Figure S4). Overall, a strong pro-tumor effect of M2 macrophages was observed on both CRC and PDAC tumor cells.

### M1/M2 macrophage-derived soluble factors mediate anti/pro-tumor effects

To explore the mechanism for the observed anti/pro-tumor effects of M1 and M2 macrophages, respectively, we collected the flow through medium from triplicate devices and subjected the medium to proteomics analysis. A total of 92 soluble factors, including biomarkers related to tumor immunity and tumor development, were included in a pre-designed immune-oncology panel (Olink Proteomics). Out of the 92 biomarkers, 50 were detectable in at least one of the samples (Table S1). Using the regular EGM-2 medium as a negative control, relative quantitation was performed between CRC663 + THP-1 M1 versus CRC663 + THP-1 M2 macrophage samples. CXCL9 (^~^ 50x), CXCL10 (^~^ 70x) and CXCL11 (^~^3x) were dramatically increased in TOC devices containing THP-1 M1 macrophages (Figure 4A). In contrast, in TOC devices containing THP-1 M2 macrophages, MMP7, ANGPT2, CCL3 and CSF-1 were significantly increased with MMP7 increased more than 7 times and ANGPT2 increased 20 times compared with the samples from THP-1 M1-TOC devices (Figure 4A).

**Figure 4:**
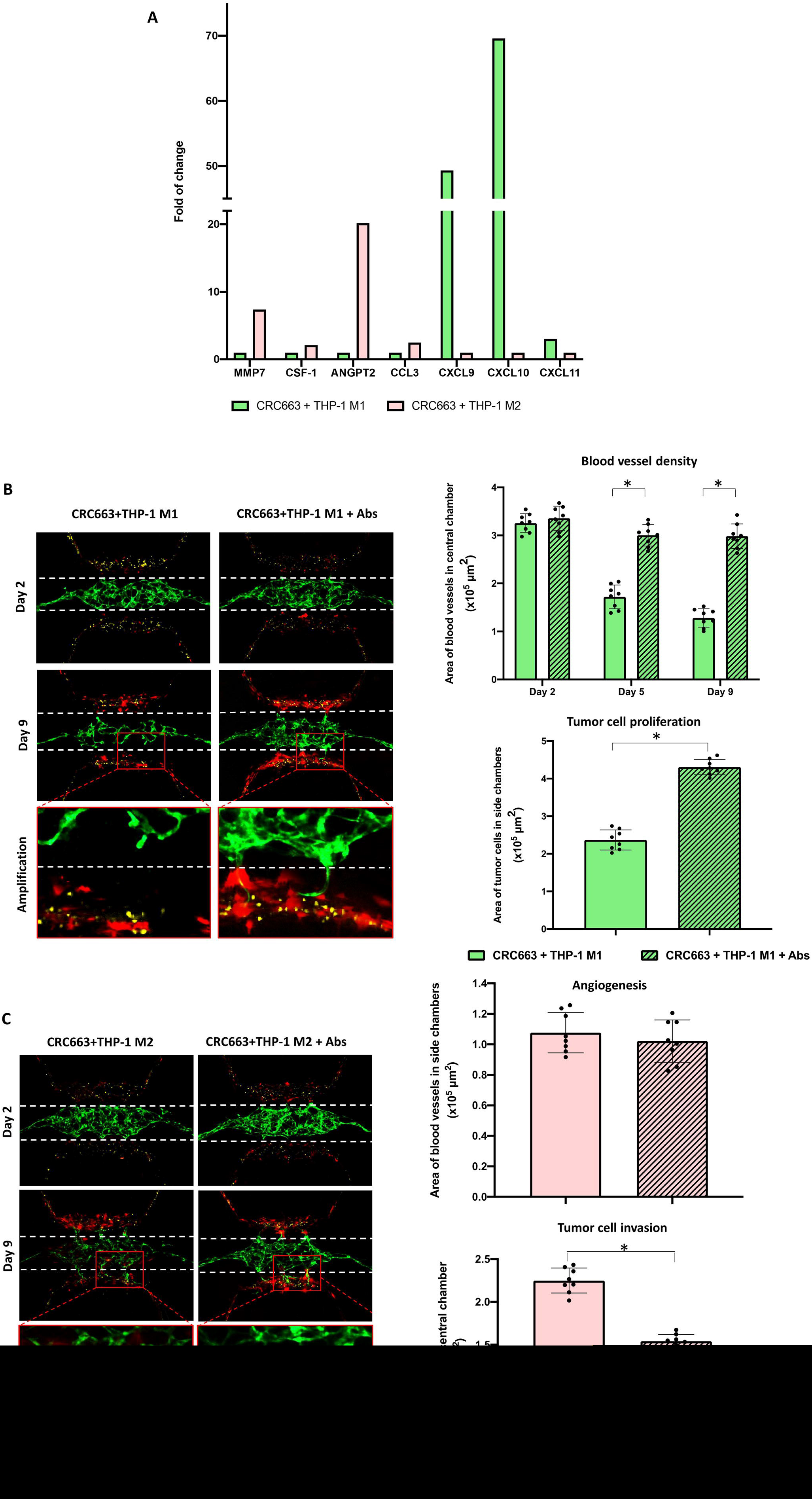
Analysis of soluble factors in M1 versus M2 macrophage-containing TOC devices show macrophage-mediated antitumor and protumor effects, respectively. A: The proteomic analysis of soluble factors recovered from TOC devices was performed. B: Antibody-mediated neutralization of chemokines in M1 TOC devices. Left panel: representative images from Days 2 and 9 of CRC663 + M1 macrophage TOC devices with or without anti-CXCL9, anti-CXCL10, and CXCL-11 neutralizing antibodies. Green: GFP labeled endothelial cells. Red: RFP labeled CRC663 tumor cells. Blue: THP-1 derived M1 macrophages. White dotted line: the top and bottom boundaries of the central chamber. Right panels: quantification of blood vessel density in the central chamber on Days 2, 5 and 9 (top panel), and tumor cell proliferation in CRC663 TOC devices with M1 macrophages on day 9 (bottom panel). C: Antibody-mediated neutralization of chemokines in M2 TOC devices. Left panel: representative images from Days 2 and 9 of CRC663 + M2 macrophage TOC devices with or without anti-MMP7 and anti-ANGPT2 neutralizing antibodies. Green: GFP labeled endothelial cells. Red: RFP labeled CRC663 tumor cells. Yellow: THP-1 derived M2 macrophages. White dotted line: the top and bottom boundaries of the central chamber. Right panels: quantification of blood vessel density in side chambers on Day 9 (top panel), and tumor cell invasion in CRC663 TOC devices with M2 macrophages on day 9 (bottom panel). Eight devices per group were used for quantification. *: p<0.05.

To test whether the differentially expressed soluble factors were directly responsible for the observed anti/pro tumor effects, we used neutralization antibodies. Mixed neutralization antibodies to CXCL9, CXCL10 and CXCL11 eliminated the anti-tumor phenotype observed in the presence of M1 macrophages (Figure 4B). For example, the microvascular network area in the central chamber of THP-1 M1 TOC devices was decreased by <10 % on day 9 compared to day 2 in the presence of the antibodies while in the untreated devices, the area of microvascular network on day 9 decreased > 50% compared to day 2. Additionally, tumor cell proliferation was restored with nearly twice as many tumor cells in the presence of antibodies (Figure 4B).

A similar reversal of phenotype was observed when neutralizing antibodies to MMP7 and ANGPT2 were added to TOC devices with THP-1 M2 macrophages (Figure 4C). Although there was no detectable change in the level of angiogenesis, the number of tumor cells that invaded the central chamber was reduced by one third in the presence of anti-MMP7 and anti-ANGPT2 antibodies (Figure 4C). Together, these data demonstrate that soluble mediators derived from macrophages including CXCL9, CXCL10, CXCL11, MMP7 and ANGPT2 contribute to the observed anti/pro-tumor effects of M1 and M2 macrophages, respectively.

### scRNA-seq identifies gene expression profiles in endothelial cells driven by tumor cells and M2 macrophages

To more deeply understand the TOC devices with higher cellular resolution, cells were harvested from devices, processed into single cell suspensions, and subjected to single cell RNA sequencing (scRNA-seq). We focused on devices with vasculature only; vasculature + tumor, and vasculature + tumor + M2 macrophages. TOC devices with THP-1 M1 macrophages were omitted due to the substantial anti-angiogenic effect (Figure 3) that would negatively impact the cell yield. As depicted in Figure 5A, all the cell types introduced in devices were recovered in the scRNA-seq data which indicates that all cell types were viable in the TOC. Within individual cell types, it appeared there were multiple clusters per cell type. When UMAP data plots of the three different types of devices were merged, the pattern of the cell clusters detected was generally similar, but did not entirely overlap, suggesting that the gene expression profiles of cells recovered from different device compositions were not the same. Introduction of tumor cells into a vascularized device triggered both upregulation and downregulation of many genes in both endothelial cells and fibroblasts (Figure S5). The addition of THP-1 M2 macrophages dramatically impacted gene expression in both endothelial cells and fibroblasts. The gene expression changes were most pronounced in the endothelial cells. For example, the presence of THP-1 M2 created a heterogeneous endothelial cell phenotype when tumor cells were also present (Figure 5B). One of the endothelial cell clusters, P2, appeared similar in all three types of devices; a second cluster (P1) was clearly distinct from the first cluster, and larger in the presence of THP-1 M2 macrophages and tumor. These data suggest that the differences between the two endothelial cell clusters were induced by tumor cells and further enhanced in the presence of pro-tumor THP-1 M2 macrophages.

**Figure 5:**
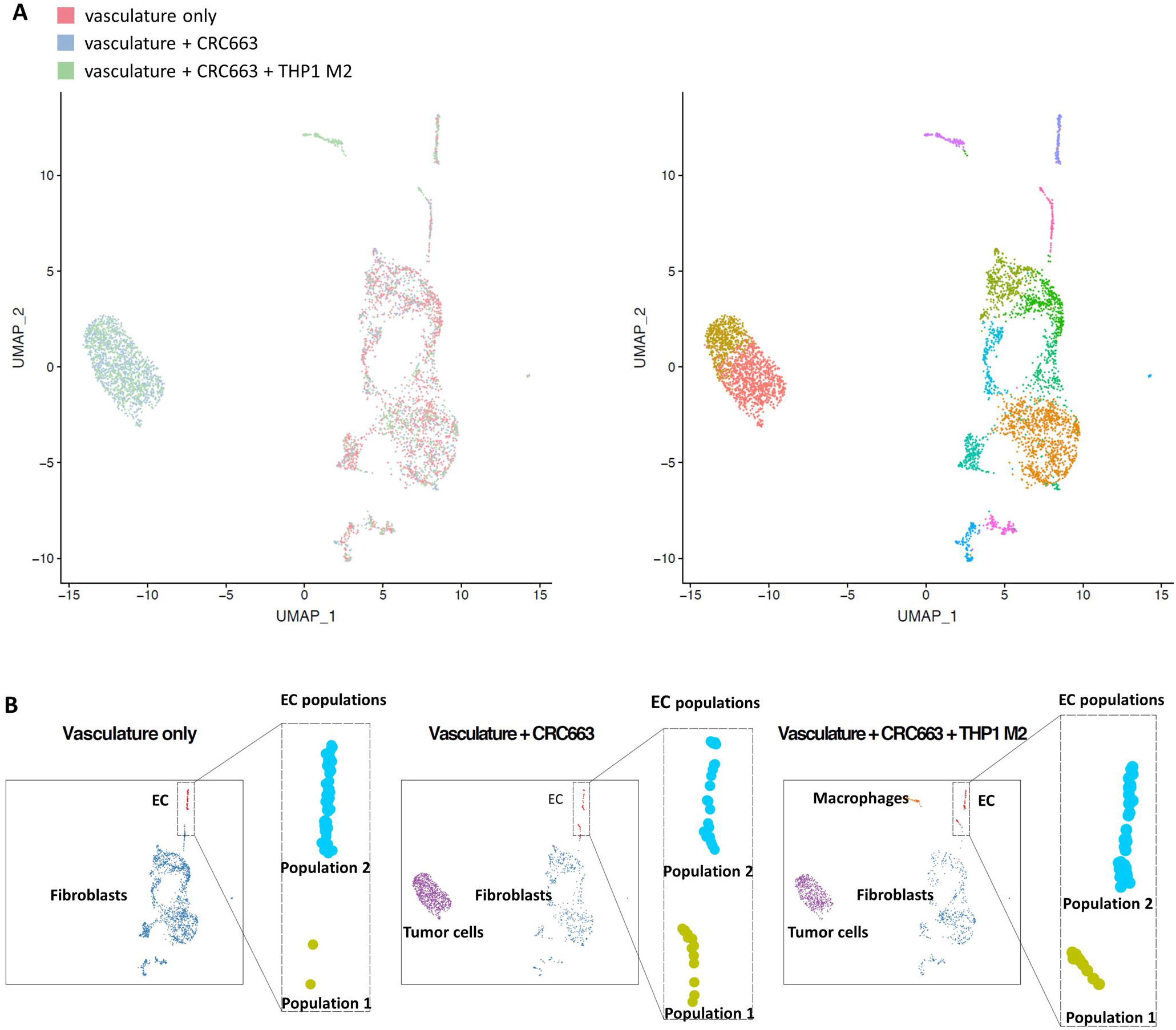
scRNA-seq analysis of TOC devices. A, left panel: integrated images of scRNA seq data of all cells recovered from three groups of TOC devices: vasculature only, vasculature + CRC663 and vasculature + CRC663 + THP-1 M2 macrophages. Right panel: integrated images of sub-clusters of cells based on the similarities of their gene expression profiles. B: normalized individual images of cells recovered from each TOC, focused on endothelial cell clusters.

### Distinct gene expression profiles in endothelial cells reflect different levels of angiogenic activity

To assess in more detail what changes tumor cells and M2 macrophages induce in endothelial cells, we compiled an abbreviated list of genes that were most differentially expressed in the two endothelial cell clusters across three different devices. A heat map was compiled of 31 genes that were most significantly changed out of > 1000 differentially expressed genes comparing the three different device conditions: vasculature only, vasculature with tumor, and vasculature with tumor and THP-1 M2 macrophages (Figure 6A). Differentially expressed genes between the P1 and P2 endothelial clusters is evident when THP-1 M2 macrophages were present (Figure 6A). Pathway analysis revealed that the genes highly expressed in endothelial cluster P1 were related to cell cycle, and immune pathways while the genes highly expressed in endothelial cluster P2 were related to tumor development and tumor invasion pathways (Figure S6).

**Figure 6:**
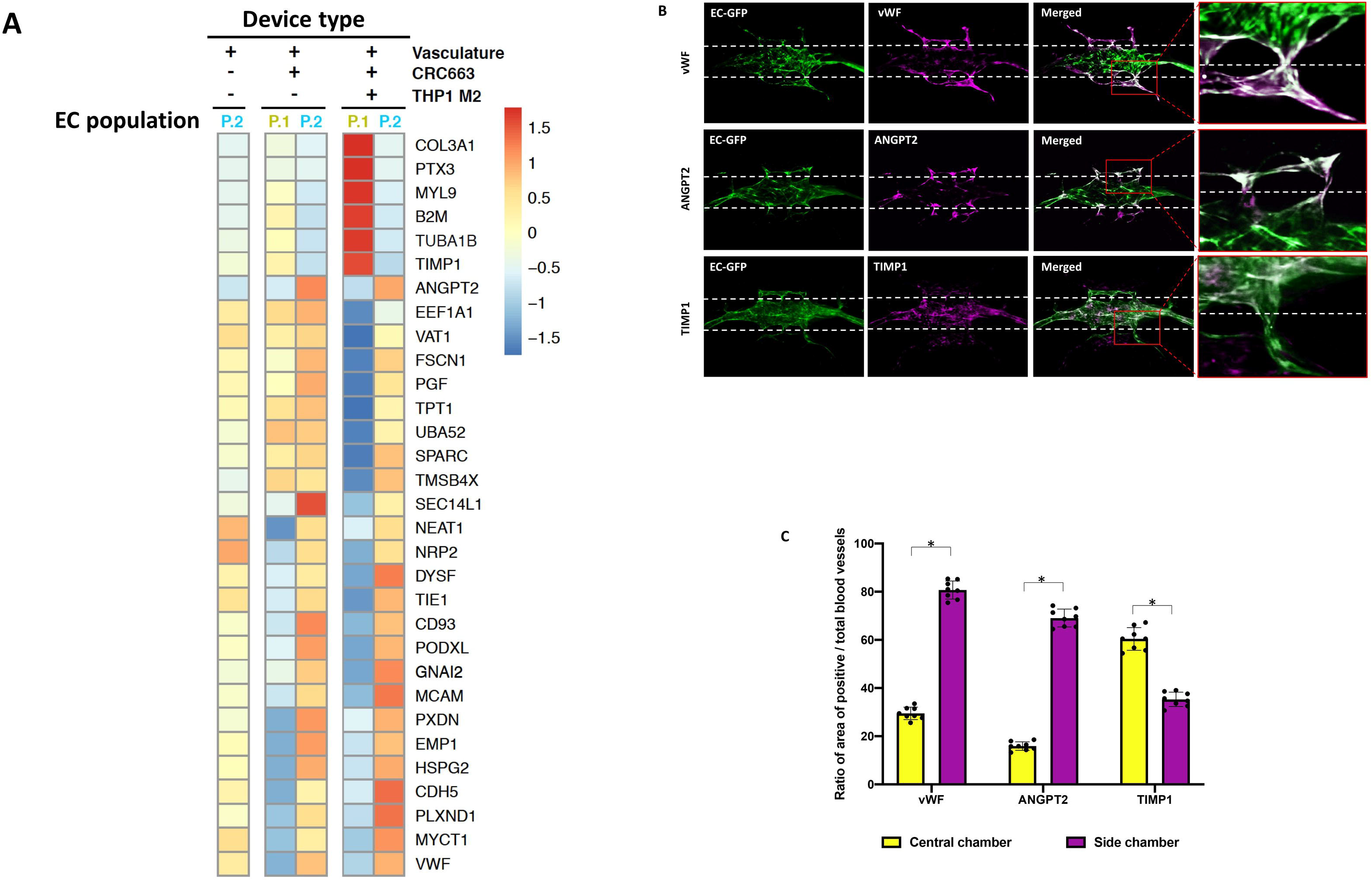
Distinct gene expression profiles in endothelial cells reflect different levels of angiogenic activity. A: Heat map of the most significantly differentially expressed genes in P1 and P2 endothelial cell clusters across all three groups. B: vWF, ANGPT2 and TIMP1 IF images of vasculature + CRC663 + THP-1 M2 devices taken on day 9. Green: GFP labeled endothelial cells. Red: RFP labelled CRC663 tumor cells. Magenta: vWF/ANGPT2/TIMP1 signal. Merged: image overlap of all three channels. C: quantification of the ratio of the vWF/ANGPT2/TIMP1 positive vasculature area (the overlapping area of magenta and green signals) versus total vasculature area (total green signals) in the central chamber and both side chambers. Eight devices were used for quantification. *: p<0.05.

We next investigated whether the endothelial cells in P1 and P2 clusters had a distinct spatial distribution within the device. We chose to analyze two genes (vWF and ANGPT2) that were highly expressed in cluster P2 and one gene (TIMP1) that was highly expressed in P1. Immunofluorescence analysis of vWF and ANGPT2 in TOC devices seeded with M2 macrophages showed that the expression of these proteins was enhanced in the microvessels present in the tumor chamber and in some areas in the central chamber where blood vessels had sprouted (Figures 6B, C). In contrast, endothelial cells with enhanced expression of TIMP1 were mainly located in the central chamber. These data suggest that the P1 and P2 endothelial clusters correspond to stable blood vessels and those engaged in active angiogenesis, respectively.

## Discussion

We recently previously reported on the design and initial validation of a novel microfluidic tumor-on-a-chip model in which tumor cells can survive and grow in a vascularized environment [7]. In Herein, we extend this model by the current report we utilized utilizing our TOC device to investigate the role of macrophages on tumor progression. Using patient-derived tumor cell lines from pancreatic adenocarcinoma and colorectal carcinoma, we discovered that macrophages with a predominant M1 phenotype create an anti-tumor microenvironment that inhibits tumor growth, migration and angiogenesis, whereas a predominant M2 phenotype creates a pro-tumor microenvironment that stimulates tumor growth, migration and angiogenesis. The anti-tumor behavior is mediated in part by chemokines CXCL9, CXCL10, CXCL11, whereas the pro-tumor behavior is mediated in part by MMP7 and ANGPT2. Using scRNA-seq, we were able to identify two distinct endothelial cell phenotypes created by the presence of the M1 and M2 macrophages in the presence of tumor cells, and map these phenotypes to regions of the tumor microenvironment that reflect stable vessels and active angiogenesis, respectively. We conclude that our TOC model is able to recapitulate important features of the tumor immune and vascular microenvironments, and can be used to understand the mechanisms by which the immune response can inhibit or stimulate tumor progression.

Tumors are complex mixtures of primarily tumor cells, leukocytes, stromal cells (e.g., cancer-associated fibroblasts (CAFs)), and blood vessels. There is ample evidence that each of these major cell types impacts tumor progression. In addition, the spatial and temporal relationships between these cells impact their function and further increase the complexity of the tumor microenvironment. It is difficult to tease apart these mechanistic relationships in small animal models, and traditionally, *in vitro* cancer modeling has relied on analysis of tumor cells grown in 2D cell culture. By their 2D and static design, these systems do not reconstitute the *in vivo* cellular microenvironment and mechanical properties found *in vivo*. The recent introduction of tumor organoid/spheroid models, including heterotypic models containing multiple types of cells, has dramatically improved our ability of to grow tumors under more natural conditions [19–21]. However, for questions related to dynamic tumor processes, such as angiogenesis, models that contain a dynamic, vascularized tumor microenvironment are required. While some groups have performed cocultures of endothelial cells and tumor cells or tumor spheres in microfluidic devices, these models do not fully recapitulate the tumor vascular network [22–26]. Our group has developed various iterations of 3D perfused vascular networks [27–29], including the current device that includes the ability to introduce and combine different cell types within the tumor microenvironment with spatial and temporal control [7]. Tumor cells and other types of cells can be introduced in two chambers that are connected to a central chamber containing an intact, pre-formed and perfused living vascular network. While most tumors readily grow inside TOC devices, our current studies also show that tumor behavior in TOC devices recapitulates important features of the parental tumors. Some tumors (CRC268, CRC663) showed enhanced angiogenesis as well as high levels of tumor migration/invasion while others showed lower levels of angiogenesis and migration. These phenotypes resembled the corresponding *in vivo* pathological features observed in the corresponding parental tumors. Other research groups have presented similar microfluidic TOC models containing perfused vasculature to study tumor-induced angiogenesis and drug delivery [30–34]. Together, these studies have demonstrated the importance of vascular network flow in evaluating tumor progression, and suggest that vascularized TOC models could be ideal *in vitro* models for personalized cancer research while complementing other *in vitro* or *in vivo* models.

We used our vascularized TOC to investigate mechanisms by which macrophages impact tumor progression. Of all immune cell subsets, macrophages, in particular the M2 phenotype, have been proven to be crucial drivers of chronic cancer-associated inflammation, are involved in almost every step of cancer progression [35, 36], and have been implicated in driving resistance to therapy [37, 38]. Tumor associated macrophages (TAMs) are associated with the M2 phenotype, and are generally recognized as potential biomarkers for diagnosis and prognosis of cancer, as well as potential therapeutic targets for cancer [39–42]. M1 macrophages on the other hand have anti-tumor properties. Pre-polarized M1 or M2 macrophages from two different monocyte cell lines (THP-1 and U-937) remained viable in our TOC devices throughout the experiments, and triggered cellular behavior in the tumor microenvironment consistent with previous reports [43, 44]. Specifically, devices with M1 macrophages showed strong anti-tumor effects such as the degradation of vasculature and reduced proliferation and migration. These observations were due, in part, to the production of CXCL9, CXCL10 and CXCL11 that can inhibit angiogenesis induced by VEGF and bFGF by interacting with a common receptor, CXCR3 on endothelial cells [45–47]. Macrophage-derived CXCL9 and CXCL10 have also been shown to be critically important for anti-tumor immune responses [9]. In contrast, devices loaded with M2 macrophages showed significant pro-tumor effects including an increase in angiogenesis and tumor migration/invasion which was associated with increased production of MMP7 and ANGPT2. MMP7 expression in colorectal cancers is increased, and has been shown to promote metastasis through degradation of the extracellular matrix and E-cadherin [14, 48, 49]. Similarly, ANGPT2 is well known to have proangiogenic and pro-tumor activity, as well as to function in resistance to anti-VEGF therapy [50–52]. Although the neutralization of these two soluble factors showed no significant difference in angiogenesis, the tumor invasion dramatically decreased in the presence of the neutralization antibodies.

Insight into the molecular mechanisms of the observed tumor phenotypes in our TOC was performed through scRNA-seq. All cell types were successfully recovered from the TOC and identified by scRNA-seq. We were able to demonstrate that introduction of tumor cells and M2 macrophages, created two endothelial cell clusters with over 1,000 differentially expressed genes. The differences in enriched pathway of the differentially expressed genes between the two endothelial populations (Figure S5) suggest further functional or biological relevance of these two populations. One endothelial cell population expressed high proangiogenic markers while the other expressed more antiangiogenic markers. In particular, vWF and ANGPT2, two well-known factors involved in endothelial cell activation and tumor angiogenesis [53–57], were highly expressed in the endothelial cell population located spatially in the tumor compartment, while TIMP1, a known antiangiogenic marker [58, 59], was expressed in the endothelial cells present mostly in the vascular chamber. The gene expression profiles and functional status of these two endothelial cell populations may allow us to better understand the relationship between inflammation and endothelial cells during tumor progression and provide potential therapeutic targets for tumor treatments.

In conclusion, our data strongly suggest that our vascularized TOC creates a tumor microenvironment that is biologically relevant and recapitulates key features of parental tumors. The TOC can be used to probe mechanisms by which the immune system impacts tumor growth. Our results show that M1 and M2 macrophages secrete distinct profiles of soluble factors that include CXCL9, CXCL10, CXCL11, MMP7, and ANGPT2 that can dramatically alter tumor cell proliferation and migration, as well as endothelial cell phenotype. The TOC model is not limited to the use of tumor cell lines [7], but can also provide insight into personalized and precision cancer treatments, including cancer immunotherapy.

## Supplementary Materials

Supplementary Figures: Figure S1-S6.

Supplementary Table: Table S1.

## Acknowledgement

This work was supported by grants from the National Cancer Institute [R21 CA223836 and P50 CA196510], the National Institutes of Health [UH3 TR00048]. We thank the Alvin J. Siteman Cancer Center at Washington University School of Medicine and Barnes-Jewish Hospital in St. Louis, MO. and the Institute of Clinical and Translational Sciences (ICTS) at Washington University in St. Louis, for the use of the Tissue Procurement Core, which provided all CRC and PDAC patients’ samples. The Siteman Cancer Center is supported in part by a National Cancer Institute Cancer Center Support Grant [P30 CA091842] and the ICTS is funded by the National Institutes of Health’s NCATS Clinical and Translational Science Award (CTSA) program grant [UL1 TR002345]. Also we thank the McDonnel Genome Institute in Washington University provided the scRNA-seq services and the Department of Pathology and Immunology in Washington University provided the flow cytometry services.

## Conflict of interest

S.C.G. has equity in Aracari Biosciences, a startup company whose core technology involves perfused human microvessels. There are no conflicts to declare for all other authors.

## Supplemental figure legends

**Figure S1: Colorectal and pancreatic commercial cell lines in TOC devices.** Two colorectal cancer (SW480 and SW620) and two pancreatic ductal adenocarcinomas (HPAC and Panc1) were introduced into the side chambers of TOC devices with pre-formed vasculature in the central chamber. A: Representative IF images are shown from Days 2 and 9 after tumor cells were introduced. Green: GFP labeled endothelial cells. Red: RFP labeled tumor cell lines. White dotted line: the top and bottom boundaries of the central chamber. B: Quantification of tumor cell and blood vessel areas in the side chambers on Days 2 and 9. Eight devices per tumor cell line were used for quantification.

**Figure S2: Work flow and flow cytometry verification of *in vitro* differentiation of THP-1 monocytes into M1 or M2 macrophages.** A: Protocol for *in vitro* polarization of THP-1 into M1 or M2 macrophages (upper panel) with the time line of preparing macrophages for loading into TOC devices (lower panel). B, left panel: Flow cytometry analysis of THP-1 derived M1 and M2 macrophages. CXCL10 and TNF-alpha expression was evaluated for M1 differentiation, and CD206 and IL-10 expression was assessed for M2 differentiation. Un-activated THP-1 cells were used as control. B, right panel: Flow cytometry of U-937 derived M1 and M2 macrophages. CXCL10 and TNF-alpha were chosen for M1 markers and CD206 and IL-10 were chosen for M2 markers. Un-activated U-937 cells were used as control.

**Figure S3: Reconstitute colorectal cancer microenvironment with CRC663 and U-937 derived macrophages showed typical anti-tumor and pro-tumor phenotypes in M1 and M2 macrophages devices respectively.** A: Representative images from Day 2, 5 and 9 after CRC663 cells with or without macrophages were loaded into the side chambers. Green: GFP labeled endothelial cells formed vasculatures. Red: RFP labelled CRC663 tumor cells. Yellow: un-activated U-937 cells or U-937 derived M1 or M2 macrophages. White dotted line: the top and bottom boundaries of central chamber. B: Quantification of angiogenesis, tumor cell proliferation and tumor invasion in the three types of TOC devices shown in A. Eight devices per group were used for quantification. *: p<0.05.

**Figure S4: Reconstitute colorectal cancer microenvironment with PDAC162 and U-937 derived macrophages showed typical anti-tumor and pro-tumor phenotypes in M1 and M2 macrophages devices respectively.** A: Representative images from Day 2, 5 and 9 after PDAC162 cells with or without macrophages were loaded into the side chambers. Green: GFP labeled endothelial cells formed vasculatures. Red: RFP labelled PDAC162 tumor cells. Yellow: un-activated U-937 cells or U-937 derived M1 or M2 macrophages. White dotted line: the top and bottom boundaries of central chamber. B: Quantification of angiogenesis, tumor cell proliferation and tumor invasion in the three types of TOC devices shown in A. Eight devices per group were used for quantification. *: p<0.05.

**Figure S5: Venn diagram for numbers of all differential expressed genes in endothelial cells and fibroblasts across the three groups of devices.** The genes expressed in vasculature only group were used as the control group. The number of genes differential expressed in other two groups compared with the control group were showed in the diagram. The number of genes in the overlapping area showed the number of common differentially expressed genes in both of the other two groups when compared with the control group.

**Figure S6: Pathway analysis of the differential expressed genes between the two endothelial cells clusters in vasculature + CRC663 + THP-1 M2 macrophage devices.** A: The most enriched pathways in the genes relatively high expressed in endothelial cluster 1. B: The most enriched pathways in the genes relatively high expressed in endothelial cluster 2.

**Table S1: Differentially produced soluble factors in TOC devices seeded with M1 – or M2-macrophages.** TOC devices were established using CRC663 tumor cells and THP-1-derived M1 or M2 macrophages. Flow through collected at day 9 was subjected to proteomics analysis using the Olink Proteomics platform. The values shown in the table represent NPX values; 1 NPX value difference equals to 2-fold change.

