## Supplementary figures for "Tumor-on-a-chip platform to interrogate the role of macrophages in tumor progression"

Supplemental Material

### Figure S1: Colorectal and pancreatic commercial cell lines in TOC devices.

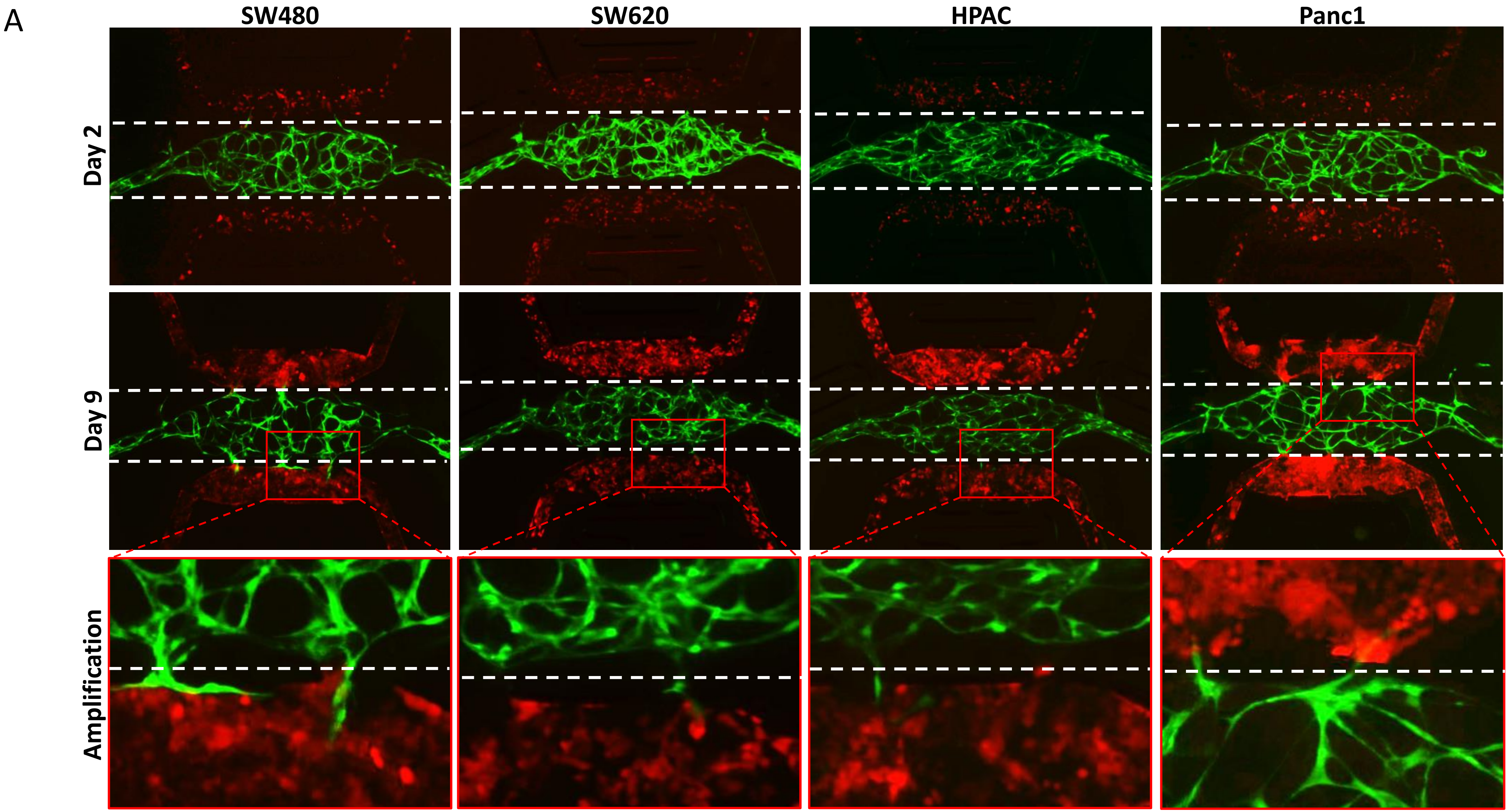

B

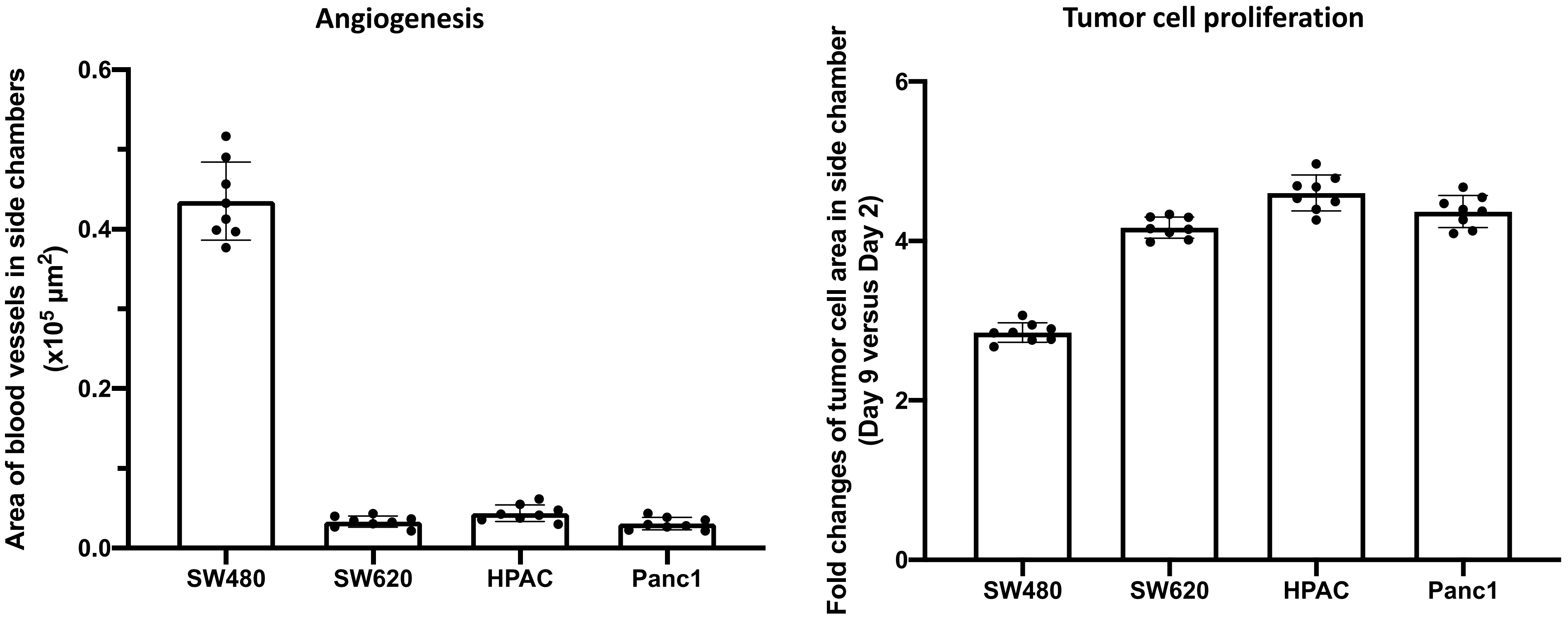

Angiogenesis
Area of blood vessels in side chambers (x10^5 μm^2)
0.6
0.4
0.2
0.0
SW480
SW620
HPAC
Panc1
Tumor cell proliferation
Fold changes of tumor cell area in side chamber (Day 9 versus Day 2)
6
4
2
0
SW480
SW620
HPAC
Panc1

#### Figure S2: Work flow and flow cytometry verification of *in vitro* differentiation of THP-1 monocytes into M1 or M2 macrophages.

**A**

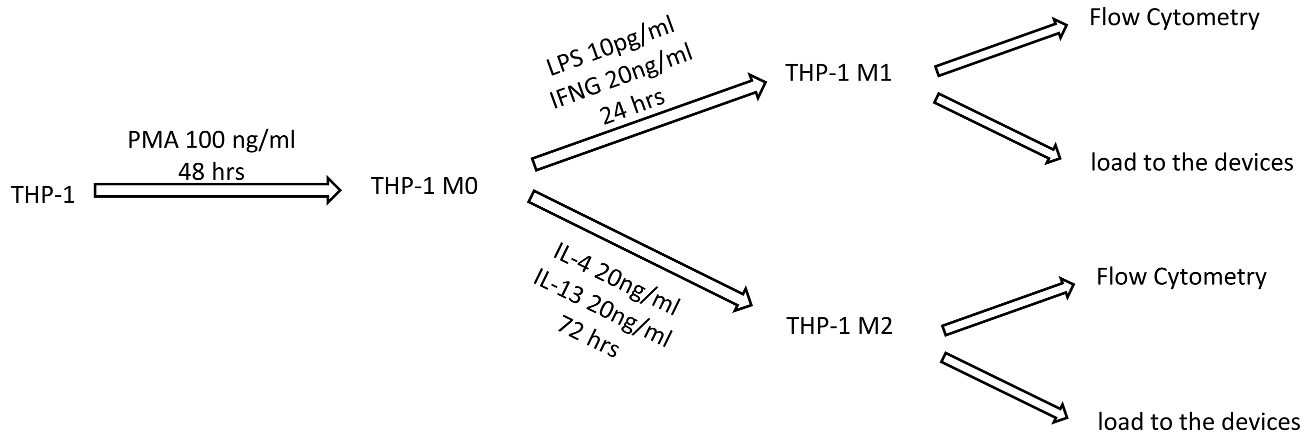


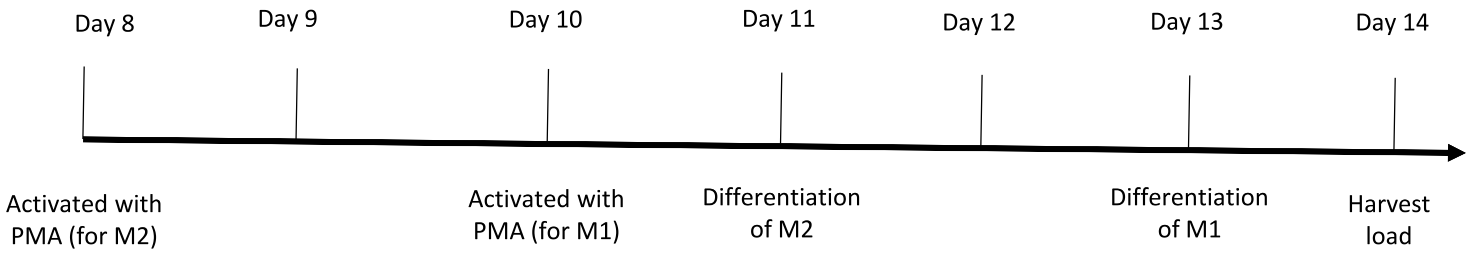

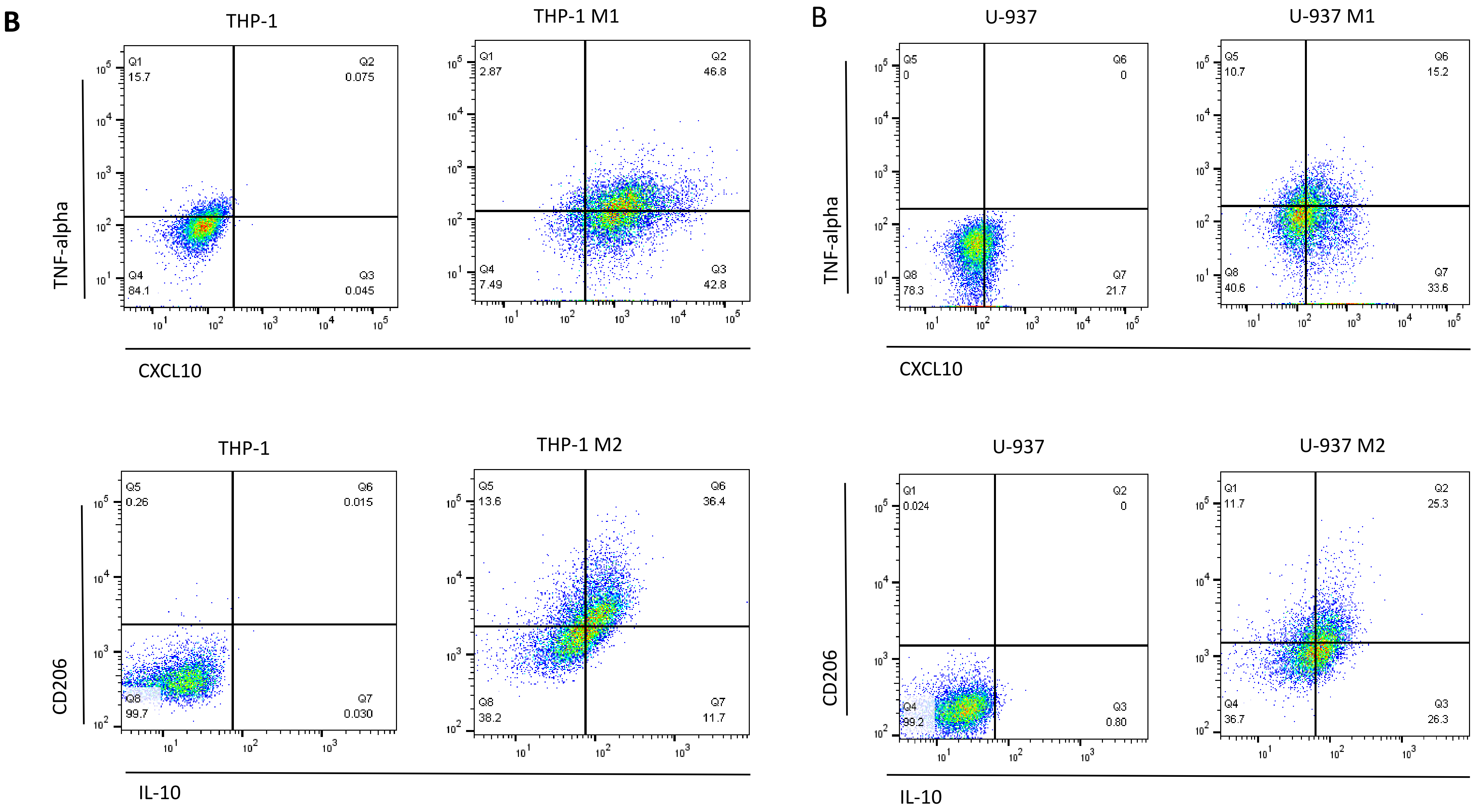

B
THP-1
THP-1 M1
TNF-alpha
CXCL10
Q1 15.7
Q2 0.075
Q3 0.045
Q4 84.1
Q1 2.87
Q2 46.8
Q3 42.8
Q4 7.49
B
U-937
U-937 M1
Q5 0
Q6 0
Q7 21.7
Q8 78.3
Q5 10.7
Q6 15.2
Q7 33.6
Q8 40.6
THP-1
THP-1 M2
CD206
IL-10
Q5 0.26
Q6 0.015
Q7 0.030
Q8 99.7
Q5 13.6
Q6 36.4
Q7 11.7
Q8 38.2
U-937
U-937 M2
Q1 0.024
Q2 0
Q3 0.80
Q4 99.2
Q1 11.7
Q2 25.3
Q3 26.3
Q4 36.7

**Figure S3: Reconstitute colorectal cancer microenvironment with CRC663 and U-937 derived macrophages showed typical anti-tumor and pro-tumor phenotypes in M1 and M2 macrophages devices respectively.**

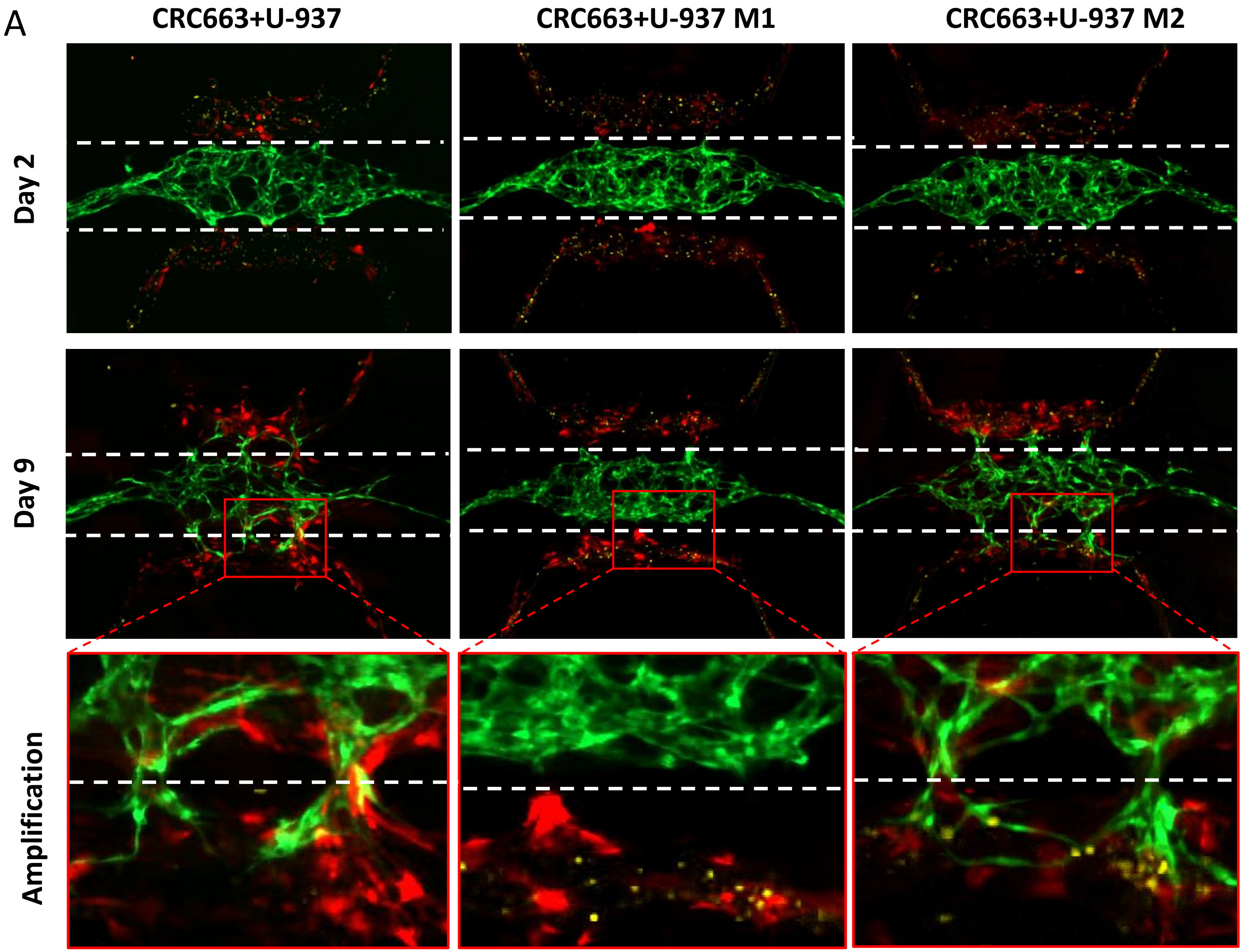

**B**

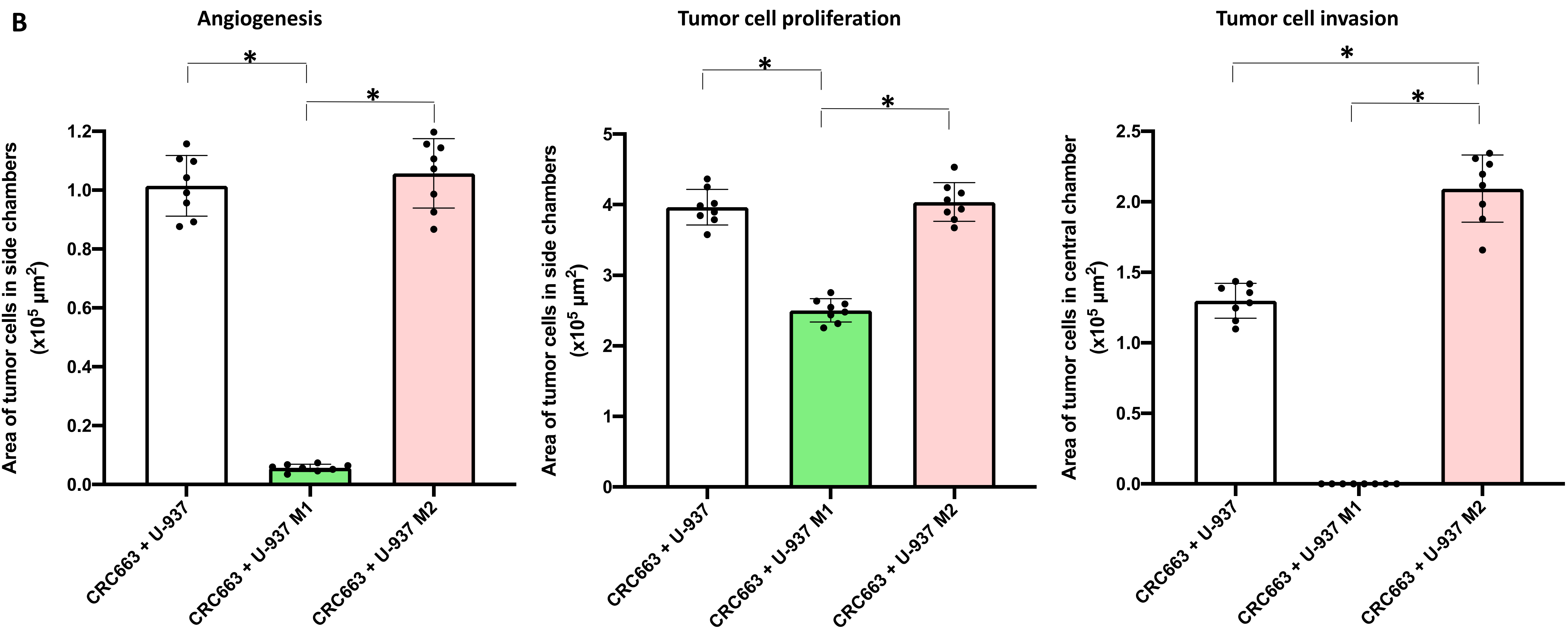
Angiogenesis
Tumor cell proliferation
Tumor cell invasion
*
*
*
*
*
*
Area of tumor cells in side chambers (x10$^5$ µm$^2$)
Area of tumor cells in central chamber (x10$^5$ µm$^2$)
0.0
0.2
0.4
0.6
0.8
1.0
1.2
0
1
2
3
4
5
0.0
0.5
1.0
1.5
2.0
2.5
CRC663 + U-937
CRC663 + U-937 M1
CRC663 + U-937 M2

**Figure S4: Reconstitute colorectal cancer microenvironment with PDAC162 and U-937 derived macrophages showed typical anti-tumor and pro-tumor phenotypes in M1 and M2 macrophages devices respectively.**

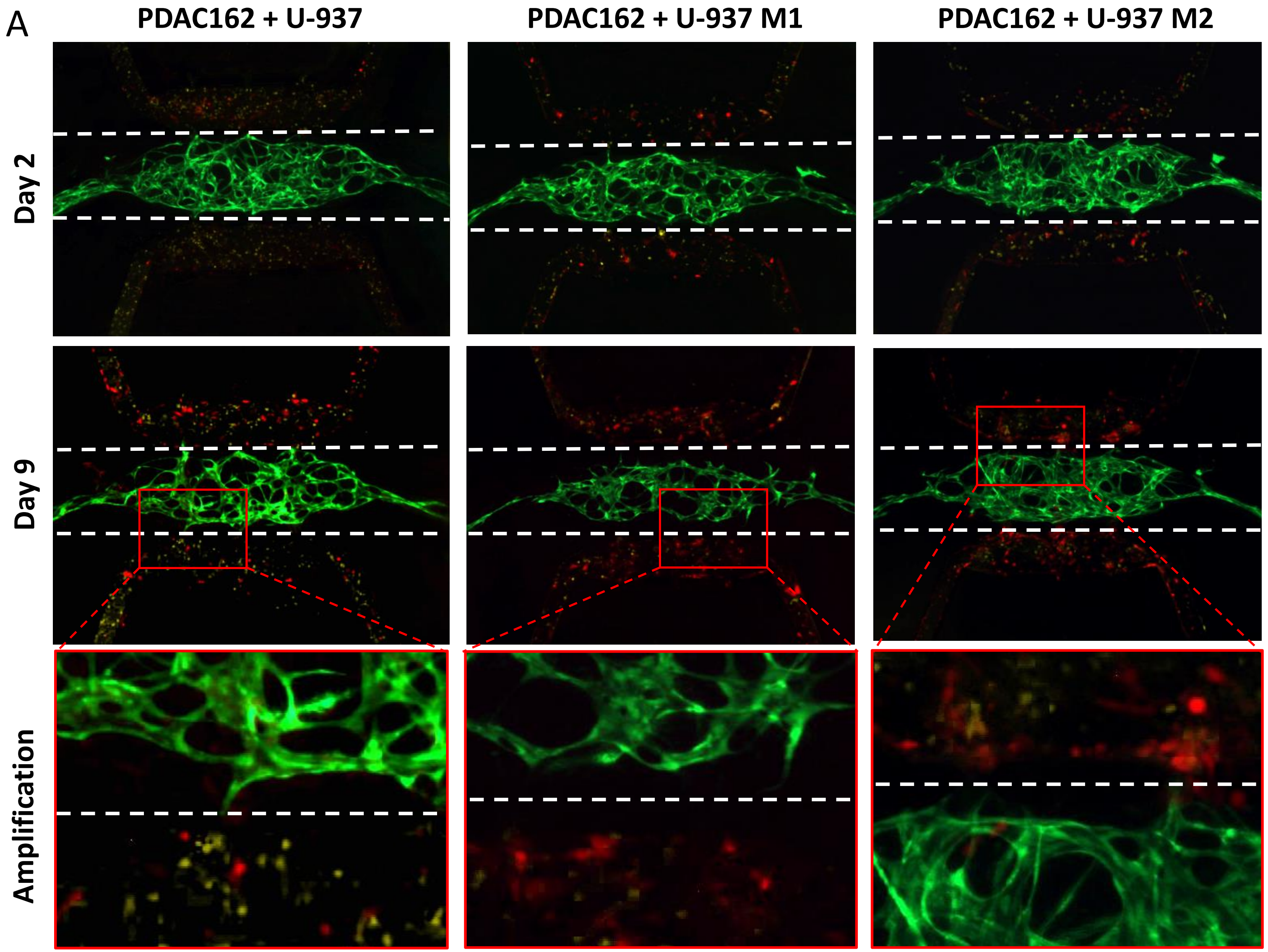

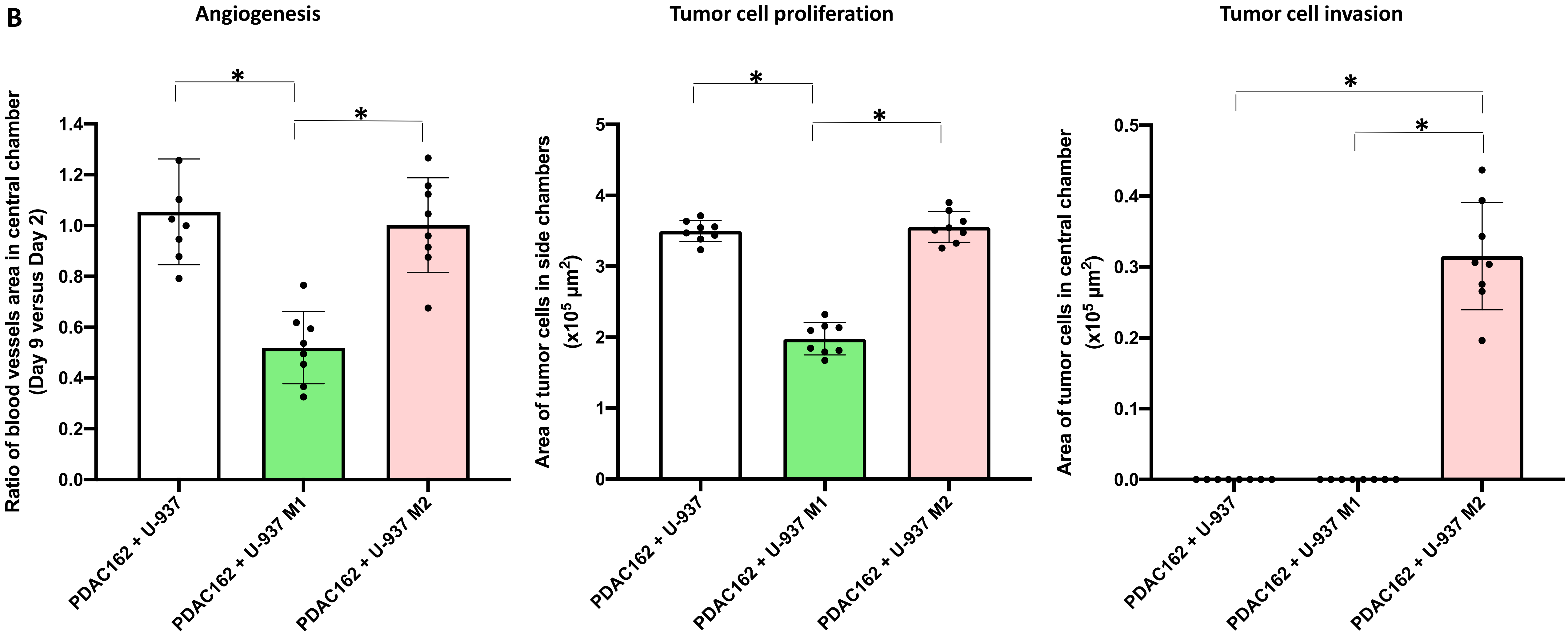

B
Angiogenesis
Tumor cell proliferation
Tumor cell invasion
Ratio of blood vessels area in central chamber (Day 9 versus Day 2)
Area of tumor cells in side chambers (x10^5 µm^2)
Area of tumor cells in central chamber (x10^5 µm^2)
PDAC162 + U-937
PDAC162 + U-937 M1
PDAC162 + U-937 M2
*

**Figure S5:** Venn diagram for numbers of all differential expressed genes in endothelial cells and fibroblasts across the three groups of devices.

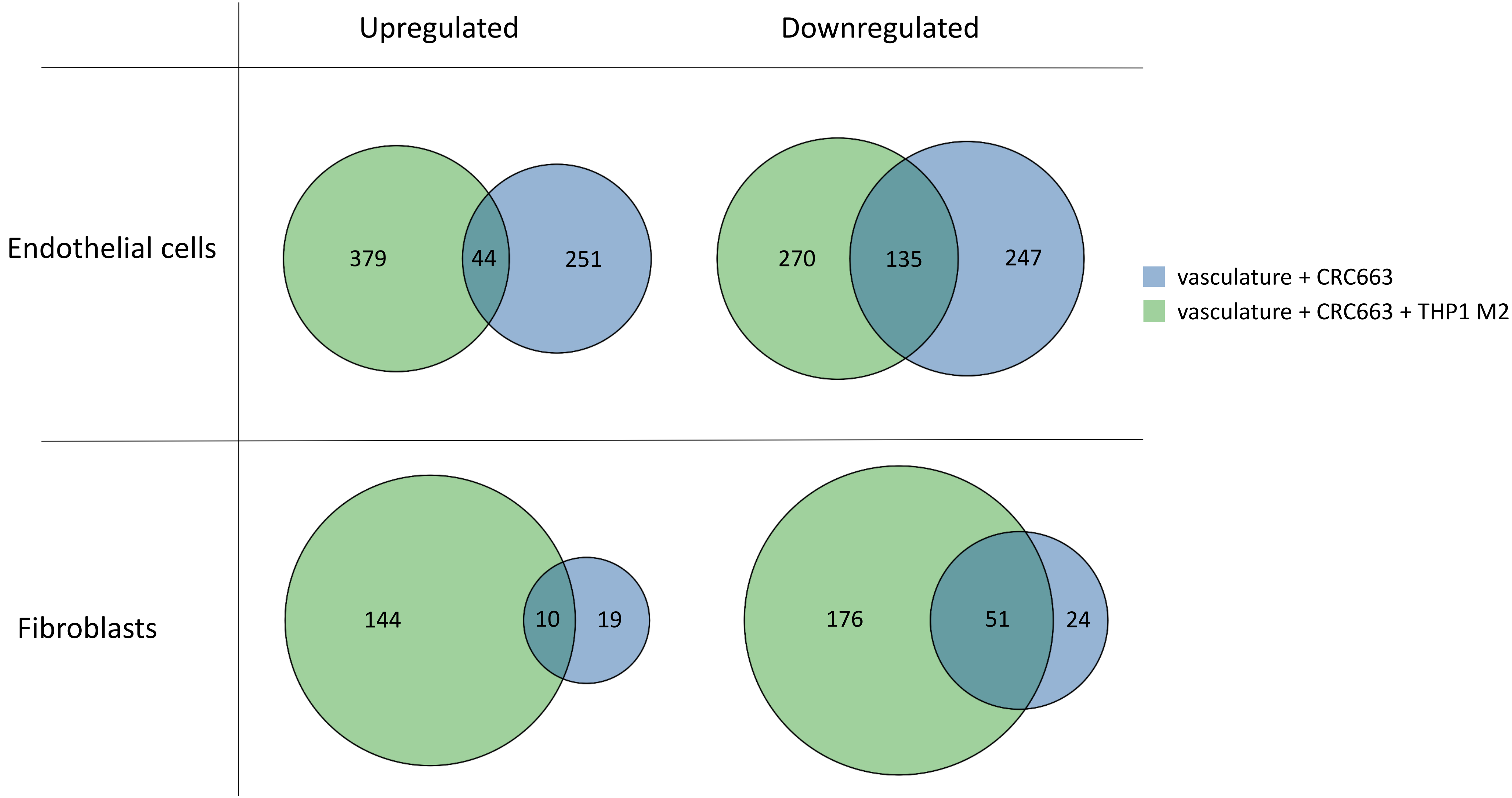

**Figure S6: Pathway analysis of the differential expressed genes between the two endothelial cells clusters in vasculature + CRC663 + THP-1 M2 macrophage devices.**

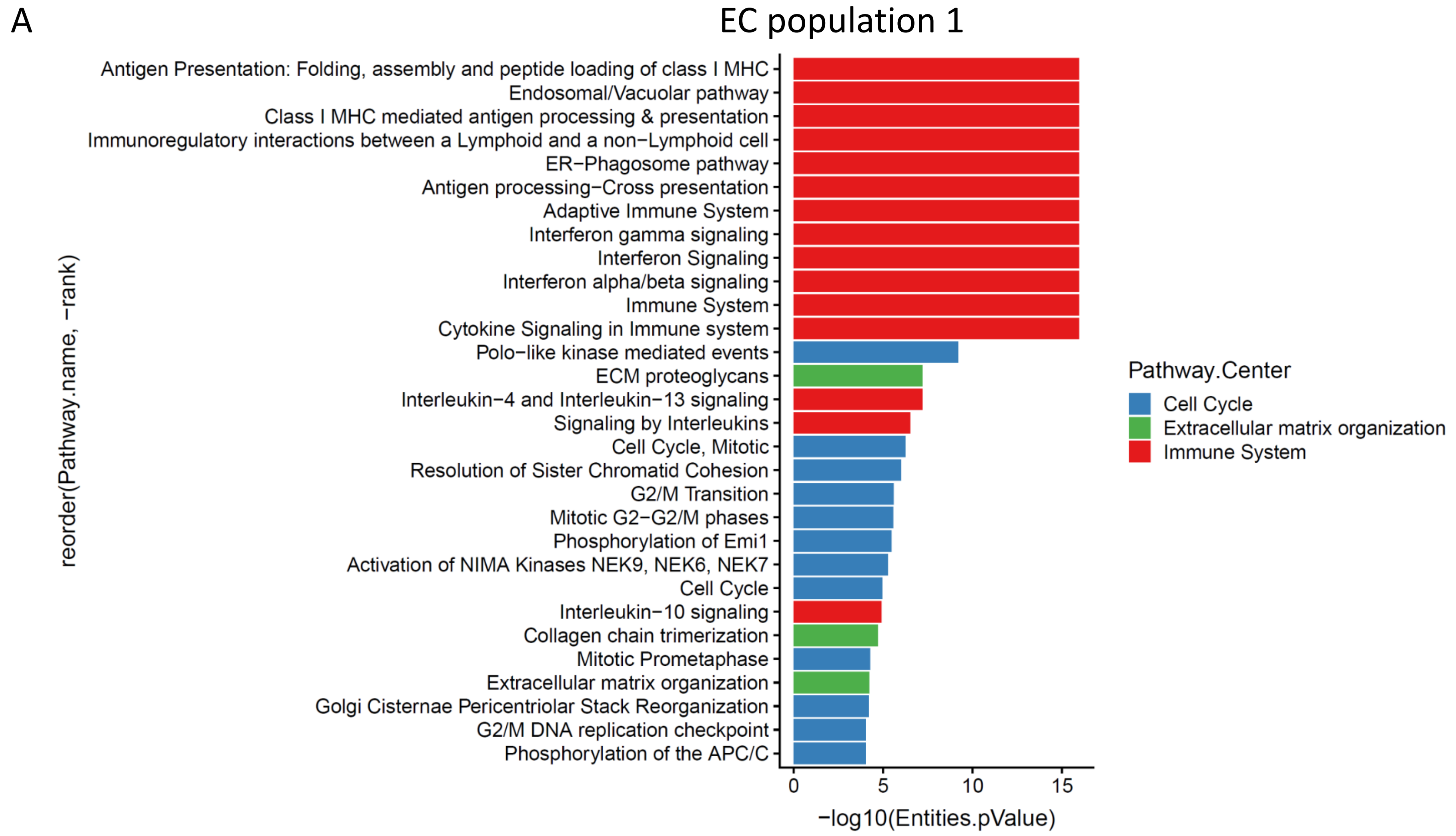

B

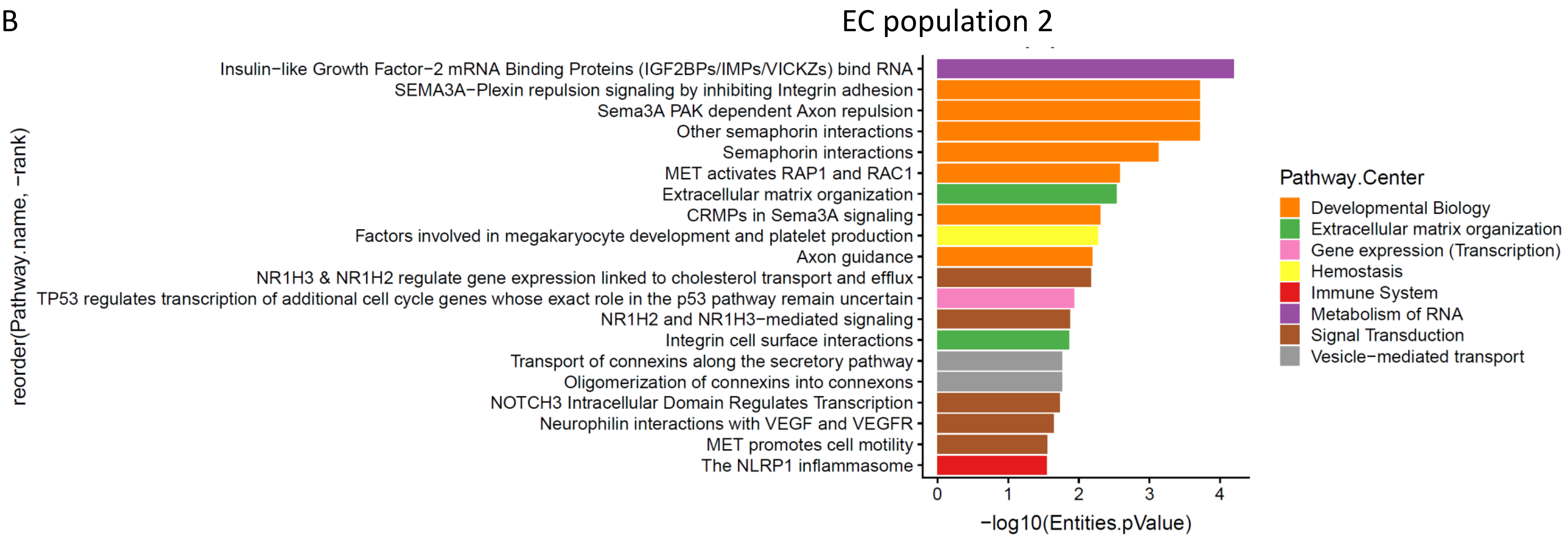

EC population 2
Insulin−like Growth Factor−2 mRNA Binding Proteins (IGF2BPs/IMPs/VICKZs) bind RNA
SEMA3A−Plexin repulsion signaling by inhibiting Integrin adhesion
Sema3A PAK dependent Axon repulsion
Other semaphorin interactions
Semaphorin interactions
MET activates RAP1 and RAC1
Extracellular matrix organization
CRMPs in Sema3A signaling
Factors involved in megakaryocyte development and platelet production
Axon guidance
NR1H3 & NR1H2 regulate gene expression linked to cholesterol transport and efflux
TP53 regulates transcription of additional cell cycle genes whose exact role in the p53 pathway remain uncertain
NR1H2 and NR1H3−mediated signaling
Integrin cell surface interactions
Transport of connexins along the secretory pathway
Oligomerization of connexins into connexons
NOTCH3 Intracellular Domain Regulates Transcription
Neurophilin interactions with VEGF and VEGFR
MET promotes cell motility
The NLRP1 inflammasome
reorder(Pathway.name, −rank)
0
1
2
3
4
−log10(Entities.pValue)
Pathway.Center
Developmental Biology
Extracellular matrix organization
Gene expression (Transcription)
Hemostasis
Immune System
Metabolism of RNA
Signal Transduction
Vesicle−mediated transport
